# In-dwelling microfluidic device for precise and reliable intranasal drug delivery during freely-moving behavior

**DOI:** 10.1101/2025.05.23.655803

**Authors:** Maria F. Ramirez, Abhishek Gour, Sarah. E. Sniffen, Emma K. Watson, Abhisheak Sharma, Daniel W. Wesson

**Affiliations:** Department of Pharmacology and Therapeutics, University of Florida College of Medicine, Florida Chemical Senses Institute, Center for Addiction Research and Education, FL 32610, USA; Department of Neuroscience, University of Florida College of Medicine, Florida Chemical Senses Institute, Center for Addiction Research and Education, FL 32610, USA; Department of Pharmaceutics, University of Florida College of Pharmacy, Gainesville, FL 32610, USA

## Abstract

Many substances/drugs are administered intranasally (IN). These include opioid overdose reversal drugs, anti-epileptic medications, migraine medications, hormone treatments, and medicines to treat/prevent allergies, colds, and flues including nasally-administered vaccines, corticosteroids, antihistamines, and decongestants. Additionally, IN administration is the preferred route of entry by users of illicit drugs. Despite the widespread use of the IN route of administration, there is no established paradigm to access this route of administration preclinically to yield precise and reliable control over delivery. This poses major gaps in therapeutic discovery/testing, establishing pharmacokinetic/pharmacodynamic relationships of therapeutics, and understanding the mechanisms of actions of therapeutics. We developed an in-dwelling microfluidic device, that, when implanted upon the nasal bone, accesses the nasal cavity to allow reliable and precise IN fluid delivery during freely-moving behavior. We validated this device, called the Nasal Access Port (NAP), to confirm it allows rapid and precise control of fluids. We further exemplified the application of the NAP for studying outcomes of IN cocaine in mice, including its pharmacokinetic profile, and both the rapid release of dopamine (DA) and behavioral effects upon IN cocaine. By achieving precise and reliable access to the IN route of administration, the NAP represents a significant methodological advance with broad applicability in the biomedical and life sciences, especially in the neuroscience, pharmacology, medicinal chemistry, and physiology domains.

## Main

Many substances/drugs are administered into the nose. These include opioids, anti-epileptic medications, migraine medications, hormone treatments, and medicines to treat/prevent allergies, colds, and flues including nasally-administered vaccines, corticosteroids, antihistamines, and decongestants^1–6^. Additionally, the nasal route of administration is the preferred route of entry by users of illicit drugs. Indeed, many drugs of abuse are inhaled by users to achieve a more immediate or intense high including cocaine, amphetamine, prescription stimulants (*e.g.,* adderall), heroin, prescription opioids, benzodiazepines, ketamine, and ecstasy^7^. Nasal inhalation of these drugs (*viz.,* “snorting”, “taking a bump”, “sniffing”, or “blowing”) holds substantial public health and treatment implications^8^, in addition to destruction and necrosis of oral, nasal, and facial structures^9–12^.

Intranasal (IN) drug administration confers multiple advantages. For one, IN delivery is ideal for drugs intended to act locally in the nasal passages, like for treatment of nasal congestion, infections, and allergies. Second, IN delivery allows for drugs to have rapid time-course of action due to the nasal cavity being highly vascularized and permeable^13^. Relatedly, drugs delivered IN have privileged access to the brain since they can bypass the blood-brain barrier – resulting in IN drug delivery being conceptually ideal for therapeutics intended to enter the brain. Indeed, inhalation of drugs into the nose exposes them to distinctive entry points whereby they can reach the CNS, including a specialized enzymatic milieu which will impact drug metabolism and unique plasma exposure. Drugs delivered IN also bypass hepatic metabolism further bolstering rapid time-course of action and bioavailability^13^. Altogether, these factors influence the total amount of drug that reaches the brain in addition to the kinetics of how it does so. IN delivery is less invasive. IN delivery has less systemic side-effects when administering therapeutics intended solely for targeting mechanisms in the nasal passages or brain^14^. And finally, from a research and development, as well as commercialization stand-point, IN delivery allows for reaching therapeutic targets and goals with less waste of valuable drugs that are otherwise metabolized and/or bind in unintended bodily compartments.

Well-developed and regulated^15^ infusion devices, designed to dispense precise and reliable doses IN, are available for testing of candidate drugs/therapies in humans. No devices are available, however, for precise and reliable IN substance administration in preclinical animal models, posing a significant barrier for early-stage therapeutic design, validation, and testing. Current preclinical methods for IN drug administration in animal models have done so by delivering the fluids using pipette tips directly onto the nares while the animal is restrained, sometimes in a supine position, and manually distributing the doses across both nostrils (*e.g.,* ^16,17^). Other methods have resorted to attaching polyurethane (PU) tubing to a syringe and delivering the fluids into the nose while the animal is anesthetized^18,19^. While these methods have highlighted the importance and viability of the IN route to deliver drugs, they do not achieve precision and reliability of drug delivery. Further, sedation and differences in handling of the animals during restraint can add variability to the accuracy with which the drug can be delivered, not to mention that this mechanism can lead to expulsion of some of the fluid due to the pressure caused by the pipette/syringe. Additionally, these mechanisms have limitations that require the experimenter to deliver higher volumes, which can lead to the fluid being inhaled into the lungs adding altering the kinetics of the nasal route. Drugs can also be vaporized or nebulized and passed into gas chambers for administration, yet here again accuracy of dosage is difficult as is it impossible to rule-out that outcomes of drugs are not due to absorption through the skin, eyes, or even mouth.

The lack of a reliable and accurate IN drug delivery paradigm presents a major void which has debilitated the drug development industry to design and test drugs which could be used to treat multitudes of medical conditions – in both human patients and companion animal patients. This void also yields uncertainty in the mechanisms of actions of drugs when delivered IN. To overcome these gaps, a paradigm is needed which achieves reliable and precise IN delivery of drugs in preclinical models.

We developed a simple-to-use surgical device, called the nasal access port (NAP), that when implanted upon the nasal bone of mice, accesses the nasal cavity to allow reliable and precise IN drug delivery during freely-moving behavior. We validated the NAP through several methods and further utilized cocaine to exemplify the application of the NAP for studying outcomes of IN drug delivery, including its rapid and precise delivery, its pharmacokinetic profile, and both the release of dopamine (DA) and robust behavioral effects upon IN cocaine delivery. This includes evidence that some animals will self-administer IN cocaine. The NAP represents a significant methodological advance with broad applicability in biomedical and life sciences research, especially in the neuroscience, pharmacology, medicinal chemistry, and physiology domains and one we predict will be a cornerstone approach for investigating IN drug delivery and its effects on genomic, molecular, cellular, systems, and behavioral consequences – all of which to-date are unexplored.

## Results

### Design and fluid dynamics of the NAP

We were inspired by prior research^20,21^ which utilized indwelling cannulae to access air pressure within the dorsal nasal recess of rodents – and worked to adapt this approach to achieve reliable IN fluid delivery. We had several criteria to reach this goal: First, the device must be able to extend into the nasal passage in a fluid-tight manner. Second, the device must be small enough to fit onto the nasal bone. Third, the device must not obstruct respiration or block the front of the nostrils, thereby allowing animals implanted with the device to freely breathe and explore their environments which rodents do prominently by protruding their snouts. Fourth, the device must have a mechanism to allow it to temporarily attach to a mating connector for fluid delivery, which can be connected and removed with little force to not risk dislodging the device from the animal. Finally, the device must have a small footprint on the cranium so to allow users requiring concurrent cranial implantation of the animals with other devices for measures and/or manipulations of physiology simultaneous with IN fluid delivery (*e.g.,* indwelling fiber optic implants for light transmittance/collection [fiber photometry and optogenetics], indwelling drug delivery cannulae, head-fixation devices, optical windows, *etc.*).

Given the importance of the size-factor in most of the above criteria, we worked to consider how to accomplish IN fluid delivery in the smallest of all main-stream preclinical animal models – the house mouse (*Mus musculus*). Like other mammals, the nasal bone covers the nasal passages which in their more posterior regions house the nasal turbinates – vascularly-rich structures which serve as the primary substrate for substances to enter the circulatory system and therefore targets of interest when administered into the nose^13^. The entire surface area of both dorsal nasal bones in an adult C57BL/6J mouse occupies only approximately 2.8mm x 6.5mm (medial-lateral x anterior-posterior)^22^. Since the nasal passages are separated by the nasal septum, fluid delivery must occur through the ventrally extending aspect of the device, which extends through the nasal bone to access a single nasal passage, covered by an even more restricted amount of cranium (approximately 1.35 mm x 6.5 mm [medial-lateral x anterior-posterior]).

With the above criteria in mind, we designed the nasal access port (NAP). The NAP is a small-footprint in-dwelling fluidic device, the body of which is constructed predominately out of the Polybutylene terephthalate (PBT) thermoplastic, Hydex®, with a 25-gauge stainless steel tube which extends throughout the NAP to achieve a channel for fluid delivery. The NAP design has four main components: 1) A connection port for fluid-tight securing of a female mating connector for subsequent fluid delivery, 2) A small magnet to achieve secure temporary fastening to the female mating connector for fluid delivery and/or a dust cap when not in use, 3) A stainless steel tube which spans the dorsal-ventral extent of the entire device, and extending beyond the base of the device, whereby the fluids can travel, and 4) An exaggerated plastic perimeter which permits for securing the NAP to the cranium. **Figure 1a** displays an image of the NAP and accompanying schematic. The largest dimensions of the NAP body span 9.9mm (anterior-posterior), approximately a third of which is devoted to the exaggerated plastic perimeter for securing to the cranium, and 10mm (dorsal-ventral), most of which is encumbered by the connection port for fluid delivery. Importantly, overall skull morphology is conserved across animals, with minimal variation in distance between cranial landmarks and skull thickness across ages, and sexes^22^.

**Figure 1.**
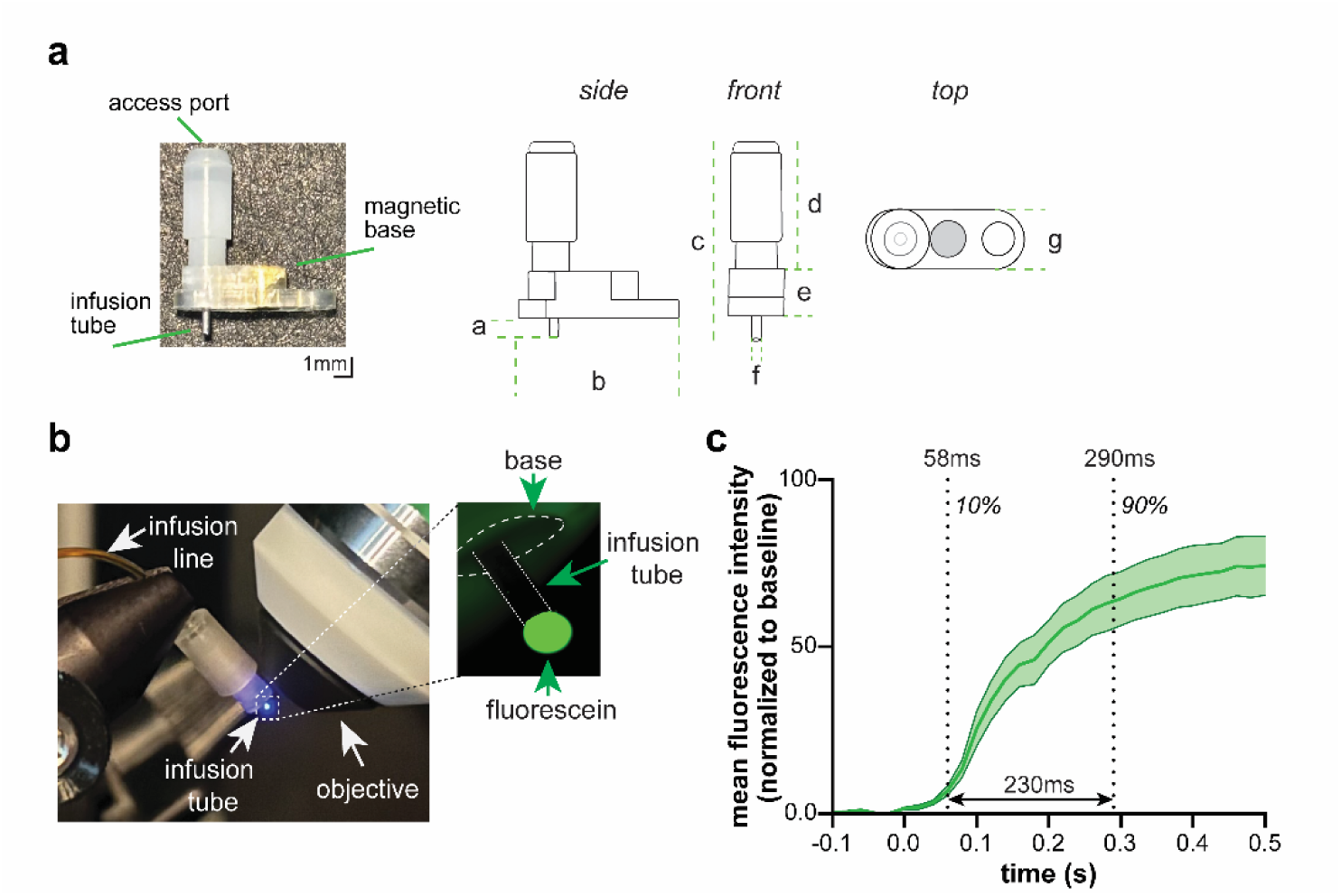
Design of the Nasal Access Port (NAP) to achieve microfluidic control. **a.** Picture of the NAP, along with schematics outlining dimensions; a=1.3mm, b=9.9mm, c=10.8mm, d=7mm, e=2.5mm, f=0.5mm, g=2.8mm. **b.** Setup for validation of fluid dynamics using time-lapse microscopy. Arrows indicate NAP attached to a flexible infusion line backfilled with fluorescein dye, with a zoomed-in inset to visualize delivery of a single 1µL droplet from the infusion port under wide-field epifluorescence microscope. **c.** Temporal dynamics of fluorescein (1µL) output from the NAP. Quantification of normalized fluorescence intensity changes from the rise (10%, 58 ± 3ms) and near-peak intensities in infusion dynamics (90%, 290 ± 19ms) across 10 trials. This quantification indicated that the vast majority of fluid is delivered from the NAP within <250ms. Data are mean ± SEM.

The selection of the 25-gauge stainless steel tube, measuring 0.5mm outer diameter, was chosen since its inner diameter (0.029mm) allows for IN delivery of small boluses of fluids with minimal resistance in order to achieve reliable flow. Importantly, the 0.5mm outer diameter fits within the medial-lateral span of a single dorsal nasal bone of the mouse (2.8mm). To record the rapid fluid-dynamics achieved by the NAP, we connected a NAP to female mating connector which was attached to a computer-controlled syringe pump via flexible PU tubing (0.43mm ID x 0.94mm OD, 25 gauge). The female mating connector and flexible infusion line were filled with the fluorescent dye, fluorescein. By positioning the NAP under a wide-field epifluorescence microscope and exciting with 455nm light, we were able to image fluid-dynamics as fluid was expunged from the NAP with temporal precision by a syringe pump (**Fig 1b**). We found that fluorescence was detected from the NAP within 58ms (± 0.003ms) of the pump being triggered (**Fig 1c**; mean of 10 replicates, 0.045ms-0.078ms across replicate means). Through identifying the rise (10%) and near-peak intensities in infusion dynamics (90%), we found that fluid boluses are released from the NAP in <250ms (**Fig 1c**; mean of 10 replicates 0.12ms-0.40ms across replicate means, 0.29±0.019ms). As in the example in **Figure 1b** (see inset), given the pump rate and tubing chosen, the boluses each exited the NAP in spherical droplets, not aerosolized. Importantly, for this purpose we chose the same syringe, pump, and approximate length of infusion line as would be appropriate for use in subsequent *in vivo* experiments therefore supporting that the NAP would yield rapid IN fluid delivery *in vivo*.

The steps for implanting the NAP for delivery of fluids into the nasal cavity are simple – following standard cranial implantation methods routinely used in physiological and behavioral neuroscience laboratories. As indicated in **Figure 2a**, the first step is to identify the mesial aspect of one of the dorsal nasal bones, particularly within the dorsal-ventral axis which corresponds to that superior to the highly vascularized nasal turbinates. In commonly used C57BL/6J mice, this is approximately 1mm anterior of the frontal-nasal fissure. Following aseptic surgical conditions, the nasal bone should be leveled by adjusting the stereotaxic incisor bar. Next, a 1mm diameter hole should be drilled through one of the nasal bones, 1mm anterior to the frontal/nasal fissure and 0.5mm lateral. The stainless-steel tube of the NAP should be lowered to extend through the craniotomy, and terminate immediately upon the nasal turbinates. Finally, the NAP should be secured with an orthopedic-compatible adhesive (*e.g.,* traditional polymer-based dental cements, UV-curable adhesives, *etc.*) to the proximal nasal bone and remainder of the skull. Following recovery, and even upon connection of mice to the infusion tether, the NAP rests upon the nasal bone in a manner which does not obstruct the nostrils nor eyes (**Fig 2b**).

**Figure 2.**
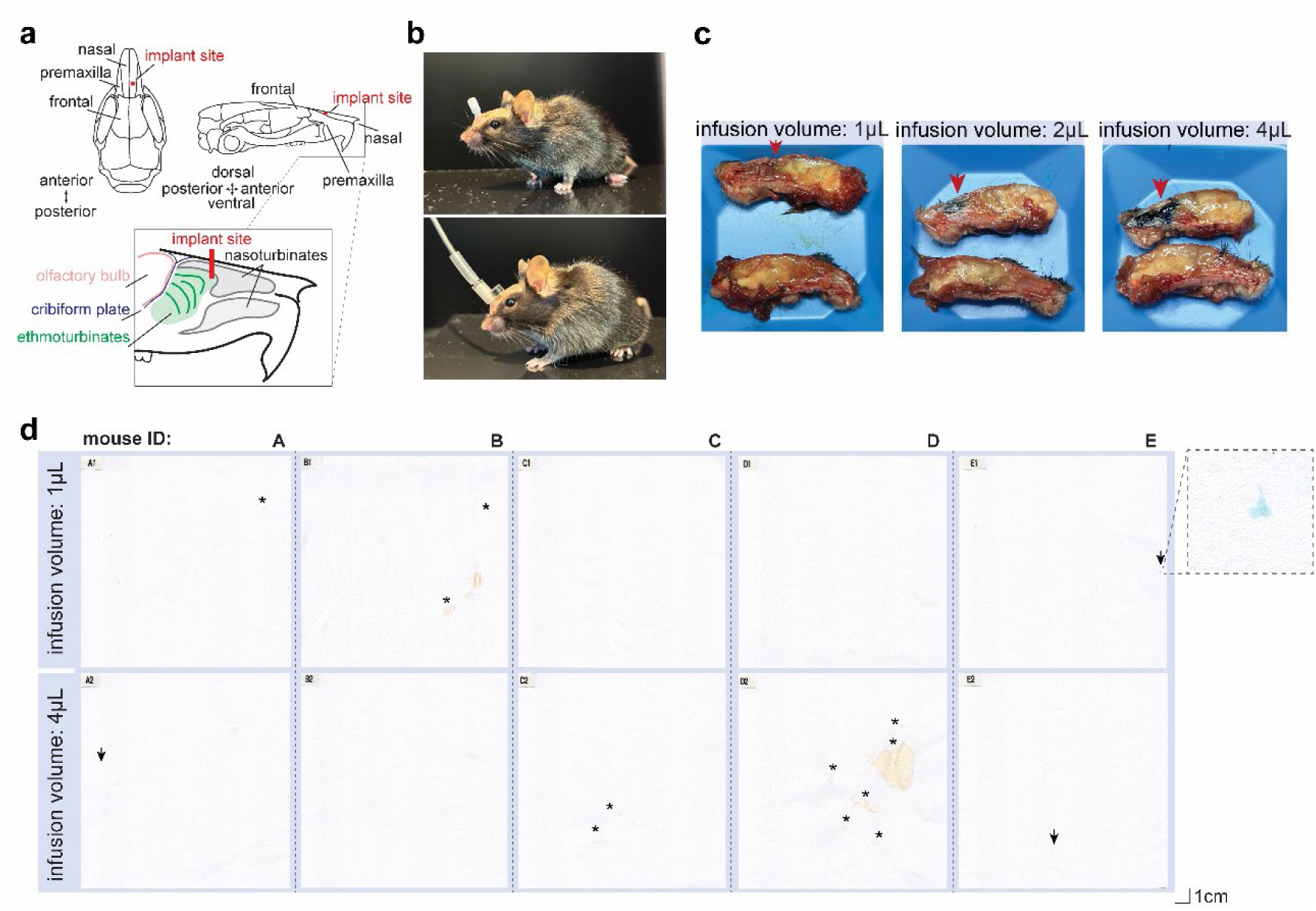
The NAP achieves precise and reliable fluid delivery which is retained in the nasal cavity. **a.** Renderings of the NAP implant site in the dorsal nasal passage of a mouse (superior view = left, lateral view = right, with inset of lateral view to illustrate approximate location of a NAP infusion tube relative to major nasal landmarks). **b.** Picture of a mouse following NAP implantation (top) and the same mouse tethered to the female mating connector attached to an infusion line within light-weight flexible spring armor (bottom). **c.** Pictures of hemisected heads to illustrate infusion spread within the ipsilateral nasal turbinates following infusion of a 1µL, 2µL, or 4µL bolus of methylene blue dye through the NAP (3 separate mice, 1/infusion volume). **d.** Images of papers, recovered from the mouse chamber floor following NAP-mediated infusion of either 1µL or 4µL boluses of methylene blue dye, with zoomed in inset (asterisks=urine, arrows=methylene blue residue; *n*=5 separate mice [labeled “A”-“E”]). Even after 4µL boluses of methylene blue dye, only trace amount of dye is detected on the papers which supports that fluids infused into the nasal cavity via the NAP are retained in the nose.

As confirmation of injection site, following anesthetic overdose, in a subset of mice we infused with 1µL of the dye methylene blue through the NAP, dissected the nasal cavity along the midline, and confirmed that methylene blue was visible on the ipsilateral turbinates following IN delivery (**Fig 2c**). Qualitatively more area of the turbinates was covered with methylene blue when larger boluses (2 or 4µL) of the dye were injected. Finally, to support utility of this approach to achieve reliable IN infusion, we investigated the possibility that IN infusion of fluids would entail escape of fluids from the nose. As above (**Fig 2c**), a subset of mice were tethered and placed into a behavioral chamber which was lined with a clean piece of white paper and subsequently infused with either with 1µL of methylene blue through the NAP, or, after changing the paper with a new clean piece, 4 rapid successions of 1µL methylene blue. As anticipated, 4 rapid successions of 1µL dye resulted in a small amount of blue dye detectable upon the papers in two of the mice, whereas the single 1µL infusion left a clean paper except trace evidence in one mouse (**Fig 2d**) – supporting that the NAP can achieve precision infusions and especially when the volume of a given bolus is kept to a minimum, fluids are retained in the nasal passages. In these same animals we also verified no detectable methylene blue in the trachea or esophagus confirming sequestration of delivery to the nasal passages.

### Application of the NAP to reliably deliver substances in vivo

Next, we sought to test the capability of the NAP for IN delivery of substances in awake, freely-moving mice. To do so, we selected to quantify known chemical read-outs of IN cocaine hydrochloride exposure. Many drugs of abuse are inhaled (*viz.,* “snorting”, “taking a bump”, “sniffing”, or “blowing”) or even applied topically to the nasal passages by users to achieve a more immediate or intense high. Exposure of drugs to the nasal surfaces holds substantial public health and treatment implications in addition to destruction of oral, nasal, and facial structures^9,11^. Despite the widespread use of the nasal route of administration by human drug users, there is no established preclinical model that accurately captures this method of administration. Cocaine is frequently administered intranasally, contributing to its reinforcing effects via recruitment of the brain’s dopaminergic system, and subsequently to its high abuse liability. However, the distinct pharmacokinetics and pharmacodynamics of IN cocaine administration in an awake mouse remain poorly understood. This knowledge gap limits our ability to develop interventions for cocaine addiction, particularly for individuals who primarily use the IN route. Further, despite the high prevalence of IN drug use, preclinical research has largely relied on intravenous, intraperitoneal, or sometimes oral routes of administration. This discrepancy limits our ability to translate findings from animal models to human drug use and hinders the development of effective prevention and treatment strategies.

To address this issue, we used the NAP to administer cocaine IN in freely-moving mice and accomplished measures of cocaine and its metabolites in brain and plasma *post-mortem*, and, in separate mice, real-time measures of DA levels in the brain utilizing *in vivo* fiber photometry^23^ with a genetically-encoded DA biosensor. Importantly, in all subsequent experiments, we validated the patency of the NAPs through infusion of a small bolus of saline and confirming that fluid was emitted from the ipsilateral nares in all mice.

NAP-implanted mice received cocaine bolus consisting of four separate 1µL infusions, achieving a cocaine dose of 8 mg/kg while they were awake and freely exploring a cage. Plasma and brain samples were collected at 1, 3, 6, or 12 min post-infusion for quantitative analysis of cocaine and its metabolites (benzoylecgonine and ecgonine methyl ester) using ultra-performance liquid chromatography-triple quadrupole mass spectrometer (UPLC-MS/MS) (**Fig 3a**). Both plasma and brain samples from treated mice contributing data to all time-points were included, as well as samples from control mice infused with physiological saline. To prevent the metabolism of cocaine by esterase, samples were stabilized with 2% potassium fluoride. IN administration of cocaine resulted in detectable levels of cocaine in the brain within just 1 min post-infusion (**Fig 3b**), with the greatest levels of cocaine in the brain (compared to saline-infused mice) were detected at 6 min post-infusion (Tukey’s multiple comparisons, *p*<0.0001) with levels started to decline by the 12 min time-point (**Fig 3b**). IN infusion of cocaine resulted in greater levels of cocaine detected in the brain compared to plasma (**Fig 3b**; two-way ANOVA, *F* (1,38) =20.89, *p<*0.0001). Consistent with its primary metabolism in the liver and plasma^24–27^, levels of benzoylecgonine, the secondary metabolite of cocaine, were detected at higher concentration in the plasma in comparison to brain (**Fig 3b**; two-way ANOVA, *F* (1,38) = 35.4, *p<*0.0001) with the greatest levels detected at 6 and 12 min post-infusion (Tukey’s multiple comparisons, 6 min *p*=0.0015; 12 min *p*<0.0001).

**Figure 3.**
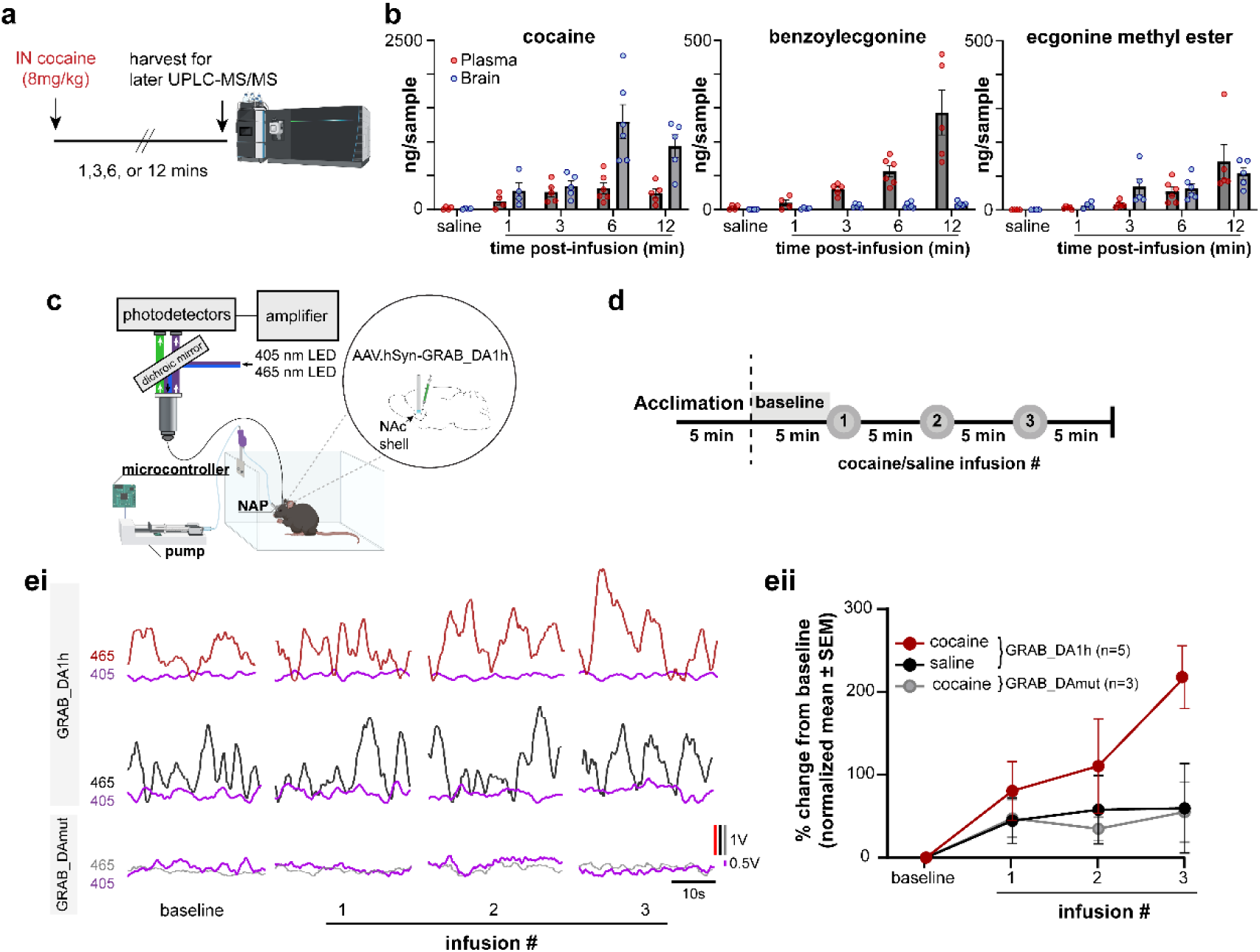
NAP-mediated intranasal delivery leads to detectable levels of cocaine and its metabolites, as well as increases in ventral striatum dopamine levels. **a.** Timeline for plasma and brain harvest post-infusion for chemical detection using ultra-performance liquid chromatography-triple quadrupole mass spectroscopy (UPLC-MS/MS). UPLC**-**MS/MS icon created in BioRender. Wesson, D. (2025) https://BioRender.com/q73a907. **b.** Plasma (ng/mL) and brain (ng/mg) levels of cocaine (two-way ANOVA, main effect of timepoint *F*(4,38)=11.99, *p*<0.0001; main effect of brain vs plasma levels *F*(1, 38) = 20.89, *p<*0.0001; Tukey’s multiple comparisons, 6min *p*<0.0001), and metabolites benzoylecgonine (two-way ANOVA, *F*(1,38)=35.4, *p<*0.0001, Tukey’s multiple comparisons brain vs plasma at 6min *p*=0.0015; 12min *p*<0.0001), and ecgonine methyl ester following infusion of cocaine (8mg/kg) or saline through the NAP at different timepoints. Data presented as mean ± SEM, points = individual mice **c.** Schematic of paradigm to monitor GRAB_DA_ signals with fiber photometry during IN cocaine deliver in freely-moving NAP-implanted mice. Some aspects created in BioRender. Wesson, D. (2025) https://BioRender.com/k25b706 **d.** Timeline of a single behavioral session, showing that following a period of acclimation, mice were delivered cocaine or saline ever five minutes. **ei.** Representative traces of GRAB_DA_ or GRAB_DA_-mut (465nm) and UV signals (405nm) in the nucleus accumbens shell (NAcSh) upon repetition of intranasal (IN) cocaine boluses (24mg/kg total dose following all three infusions). As expected, dopamine transients were detected in either saline or cocaine-infused mice at baseline (far left black and red traces, respectively), and these transients became larger following deliveries of cocaine (top right red trace) **eii.** Averaged normalized percent change from baseline of GRAB_DA_ and GRAB_DA_ -mut (grey) upon bolus repetitions of 8mg/kg cocaine (red) or saline (black) (rmANOVA *F*(1.566, 6.263)=5.059, *p*=0.054, Tukey’s multiple comparisons baseline vs. cocaine bolus 3 in GRAB_DA_ mice *p*=0.015, bolus 3 GRAB_DA_ vs. GRAB_DA_-mut mice *p*=0.024). Data are mean ± SEM.

Next, we used real-time measures to examine neurochemical outcomes following IN cocaine. Cocaine exerts its effects on neural circuitry and behavior primarily by inhibiting the reuptake of biogenic amines, including DA in a key “brain reward” center, the nucleus accumbens ^28–30^. To monitor possible changes in DA levels upon IN cocaine delivery, we injected mice with the DA sensor GRABDA (AAV9.hSyn-GRAB_DA1h)^31^ into their nucleus accumbens shell (NAcSh), and subsequently implanted an optical fiber in the same region for subsequent fiber photometry-based monitoring of DA in freely-moving mice (**Fig 3c**). All mice were also implanted with a NAP as described above. On separate days, mice were IN delivered boluses of either physiological saline or cocaine hydrochloride in a counterbalanced manner. 4 boluses were delivered (2mg/kg each, 1µL each infusion, 15sec inter-infusion interval), over three total repetitions separated in time by approximately 5 minutes, spanning 15 minutes total, yielding a cumulative amount of 24mg/kg cocaine/mouse. As shown in **Figures 3ei and 3eii**, we observed elevations in the GRABDA signal upon the second and especially the third cocaine bolus – indicating that IN cocaine via the NAP results in elevations in CNS DA levels, at least as detected by a biosensor, within <15 minutes. Increases in nucleus accumbens DA levels were detected across repetitions of cocaine boluses in GRABDA mice (**Fig 3eii**; rmANOVA, *F*(1.566, 6.263)=5.059, *p*=0.054), with the greatest increase in DA levels after the third bolus (Tukey’s multiple comparisons, baseline vs bolus 3, *p*=0.015). This increase was specific to cocaine and DA in that no such increase was detected in the same mice on the day they were administered IN saline (Tukey’s multiple comparisons bolus 3 vs saline, *p*=0.046), nor in separate mice injected with IN cocaine, yet monitored by means of a GRAB sensor mutated to be unresponsive to DA, GRABDA-mut (**Fig 3eii**, Tukey’s multiple comparisons bolus 3 of cocaine in GRABDA vs GRABDA-mut mice *p*=0.024; see Methods). Altogether, these results indicate that cocaine is detected soon-after, as are brain DA levels elevated, following IN infusion via the NAP in freely-behaving mice.

### IN cocaine rapidly and robustly drives behavioral changes

Having established that the NAP achieves rapid accumulation of cocaine in both plasma and brain, as well as DA in the brain following IN cocaine delivery, we next sought to examine if the NAP permits drug delivery concurrent with behavioral monitoring of drug-induced behavioral changes. As a starting point, we monitored locomotion in mice to identify that IN cocaine rapidly drives hyperlocomotion. NAP implanted mice were placed in chambers with infrared cameras for later tracking of distance traveled^32^ and connected to a flexible infusion line attached to an infusion pump (**Fig 4ai**). Each day, following 15 minutes of acclimation to the chamber tethered to an infusion line, the mice received a single bolus (1µL infusion, 4X, every 15sec) of saline or cocaine (**Fig 4aii**), with increasing concentrations to achieve increasing doses of cocaine across separate days. Across sexes, we found a significant increase in locomotion in cocaine-treated mice at the 4mg/kg per infusion dose, which escalated with increasing doses on subsequent days (**Fig 4b**; (mixed-effects model, *F*(4.597, 88.59)=6.998, *p*<0.001); Tukey’s multiple comparisons 4mg/kg/inf vs. saline *p*=0.024). Increases in locomotion were observed in both sexes (**Fig 4c**). In a subset of mice used above, we allowed cocaine to wash-out for at least one week and then tested the time-course whereby IN cocaine would generate hyperlocomotion by analyzing locomotor changes upon delivery of a highly-concentrated cocaine bolus (48mg/kg). This uncovered that significant elevations in locomotion occur within just 5 minutes post-infusion of IN cocaine (**Fig 4d**; two-way rmANOVA, *F*(4.552, 81.94)=6.984, *p*<0.001; Tukey’s multiple comparisons cocaine treatment vs saline at 5min *p*=0.048).

**Figure 4.**
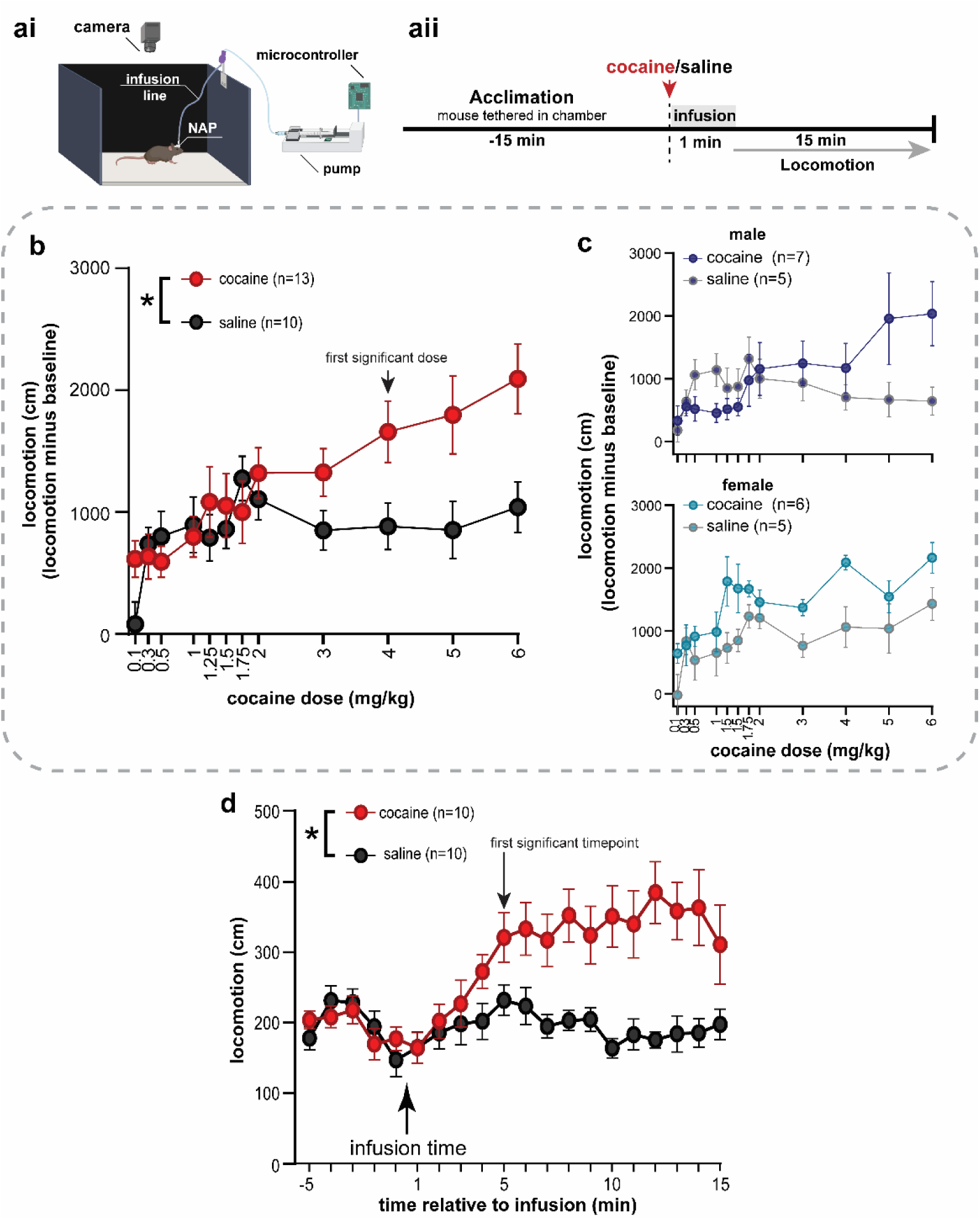
Monitoring unstructured behavioral changes following NAP-mediated IN drug delivery. ai. Experimental set-up for monitoring locomotion in mice during IN infusion of cocaine. Cocaine was automatically infused into the nose via a syringe pump controlled by a microcontroller while locomotion was recorded by an overhead camera. Some aspects created in BioRender. Wesson, D. (2025) https://BioRender.com/h63s104. **aii.** Timeline of a single behavioral session, showing that following a period of acclimation, mice were delivered cocaine or saline (separate mice / treatment) over the course of one minute, and locomotion monitored for the subsequent 15 minutes. **b.** Changes in locomotion following intranasal (IN) infusions of gradually increasing doses of cocaine or saline across consecutive days (mixed-effects model, *F*(4.597, 88.59)=6.998, *p<*0.001) with significant differences at 4mg/kg/inf (16mg/kg) of cocaine vs saline (Tukey’s multiple comparisons, *p*=0.024). Data (mean ± SEM) normalized to baseline. **c.** Sama data as in **b**, but aggregated by sex to indicate that cocaine-infused mice of both sexes showed greater locomotion than separate mice infused with saline. **d.** Time-course of hyperlocomotion evoked by IN infusion of a highly-concentrated 48mg/kg cocaine bolus (two-way rmANOVA, *F*(4.552, 81.94)=6.984, *p*<0.001; Tukey’s multiple comparisons cocaine treatment vs saline at 5min *p*=0.048).

### Validation of the NAP for use during instrumental behaviors

Intravenous self-administration models wherein mice seek and take cocaine using instrumental procedures are widely used to understand the behavioral outcomes of cocaine, to define neuropharmacological mechanisms of cocaine, and have provided insights into cocaine-induced plasticity and circuitry that differs from passive cocaine administration^33–36^. Through these paradigms models have been established, wherein like human users, some rodents will show ‘high responding’ to take cocaine, whereas others maintain ‘low responding’^37–41^. Due to these factors, cocaine self-administration paradigms are thought to have high face-validity for recapitulating cocaine use disorder in humans. Having established the NAP, we sought to test the applicability of the NAP for use in drug self-administration. Would mice with an implant on their nasal bone engage in an operant behavioral assay? Will some mice escalate in cocaine taking across sessions as reported in intravenous paradigms? Further, since irritation induced by IN cocaine itself would provide a cue to the animal as an IN stimulus, will mice work to take cocaine IN in the absence of drug-paired or contextual cues which are otherwise needed for animals to develop self-administration in intravenous paradigms^42–45^?

To test these questions, mice of both sexes were placed into operant chambers containing nose-poke ports equipped with infrared photodiodes that signal port entry and NAP’s connected to flexible infusion tethers attached to infusion pumps to develop IN cocaine (**Fig 5a**; 0.3mg/kg/ infusion, infusion of 1µL volume delivered over a 350ms period). Before the first self-administration session, active (cocaine-delivering) and inactive ports (no outcome, monitored to assay poking behavior independent of achieving IN cocaine) were randomly assigned for each chamber and mice allowed to self-administer for up to 2 hours daily (**Fig 5b**). NAP patency was verified pre- and post-testing by IN saline resulting in fluid exit from the ipsilateral nares. Consistent with prior intravenous work, we found that some mice escalated in their taking of IN cocaine across behavioral sessions. Across all mice, 27% (4/15) displayed high-responding behaviors as defined by >70% of pokes occurring in the cocaine-paired port for at least 2 consecutive sessions (**Fig 5ci)**. Among these mice, they displayed on average 85% of pokes for the cocaine-paired port compared to the inactive port on their last three behavioral sessions (**Fig 5cii**; 70-97% inter-animal range). High-responding mice displayed significant escalation in poking for the cocaine-paired port across sessions (**Fig 5cii**; two-way rmANOVA, *F*(2,12)=11.99, *p*=0.001).

**Figure 5.**
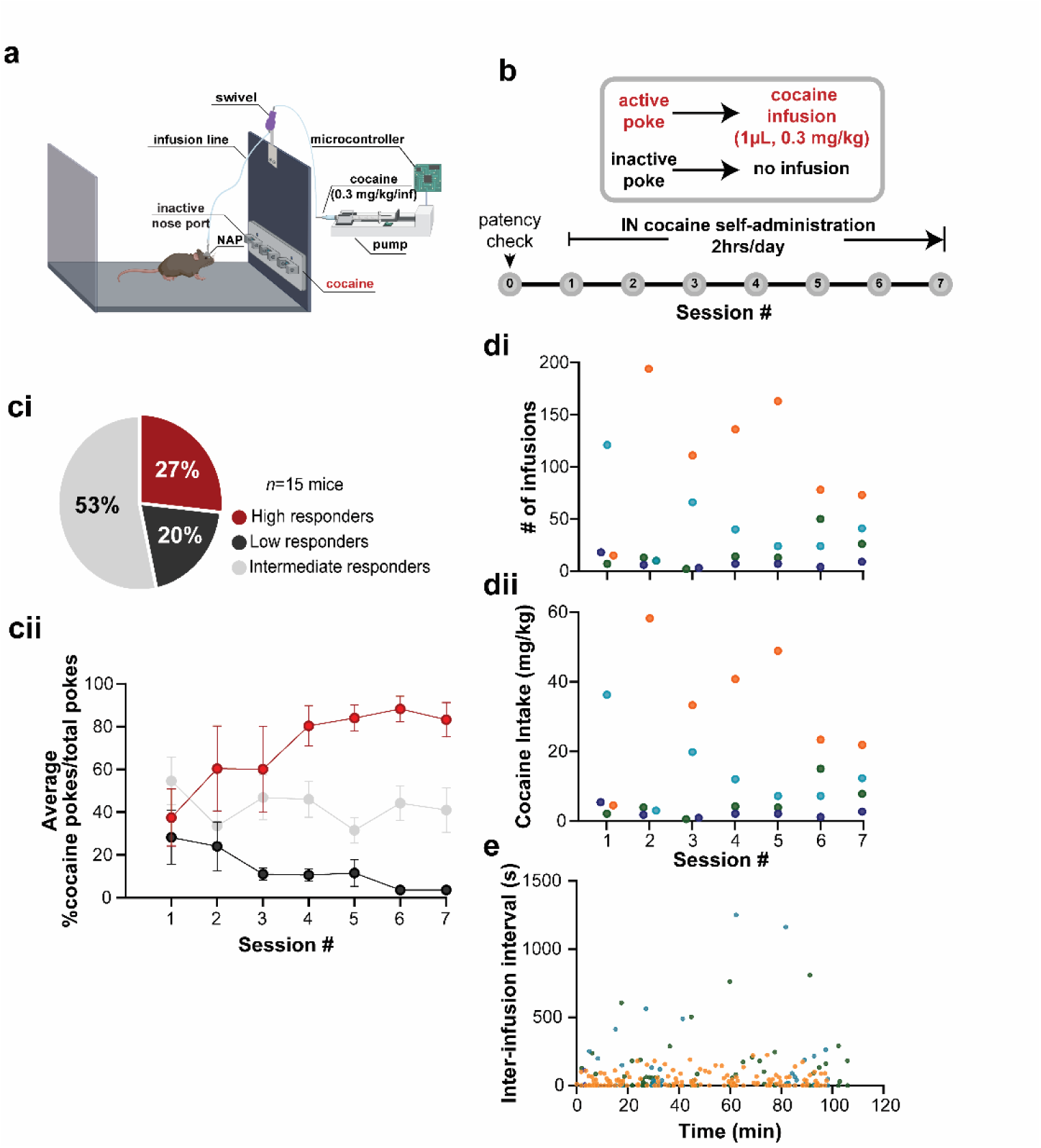
The NAP is applicable for instrumental procedures, including cocaine self-administration. **a.** Schematic of operant chamber for IN self-administration of cocaine. Some aspects created in BioRender. Wesson, D. (2025) https://BioRender.com/t32b008. **b.** Experimental timeline for behavioral sessions wherein nose-poke into an active port triggered a single infusion of cocaine IN, whereas poke into the inactive port resulted in no outcome. Cocaine infusion was controlled by a microcontroller which monitored nose-pokes and triggered a syringe pump upon active pokes. No cues other than IN delivery accompanied active pokes. **ci.** Population distribution of high-responding, intermediate-responding, and low-responding mice to IN cocaine (0.3mg/kg/inf of cocaine; infusion of 1µL volume delivered over a 350ms period). **cii.** Average percent of cocaine pokes/total pokes across 7 days displayed by high-responding, intermediate-responding, and low-responding mice. High-responding mice, as a group, displayed progressively more active pokes for IN cocaine across days (two-way rmANOVA, *F*(2,12)=11.99, *p=*0.001). Data reported as mean ± SEM. Data in **d** are from same mice as in c, to report (**di)** the number of IN infusions taken, and (**dii**) cocaine intake in mg/kg by high-responding mice across the 7 days of self-administration sessions [individual mouse data are shown by distinctly-colored points]. **e.** Inter-infusion interval exhibited by high-responding mice (same as in **di** and **dii**) within the session with the greatest number of pokes (average of 67sec, 14-119s inter-animal range). Individual mouse data shown in colors matched to **di** and **dii.**

As expected from prior work investigating cocaine-taking in intravenous rodent models^46–51^, population differences in drug-responding vulnerability also manifested from IN cocaine self-administration, which we have defined here as intermediate- and low-responding mice (those not displaying bias towards poking in the active port across sessions and those displaying decreases in poking in the active port across sessions, respectively; **Figs 5ci & 5cii**). Further validating IN cocaine self-administration was achieved in some mice, we inspected behaviors of high-responding mice to uncover that several of the mice took infusion boluses at respectively high numbers (**Fig 5di**) achieving considerably large doses of cocaine intake within sessions (**Fig 5dii**), and further largely displayed consistent motivation within individual sessions as indicated by an across mouse inter-infusion interval average of 67 sec (**Fig 5e**; 14-119s inter-animal range within the session with the greatest number of pokes).

### Consistency of in vivo fluid delivery

A common issue which plagues intravenous jugular catheter drug delivery in preclinical models is that the catheters suffer from mechanism failure or become clogged / lose patency for fluid entry into the body^52–54^. From mice throughout all the above *in vivo* experiments, we verified patency upon end of the experiments by means of either validating fluid emission from the ipsilateral nares, or by means of observing methylene blue staining of the nasal turbinates. Mice survived for various durations post NAP implant depending upon the experimental needs required of each mouse. Across all experiments, including even the behavioral assays which involved daily connecting and disconnecting of the animal to infusion lines for weeks, the NAP yielded a >91% success in maintaining patency for an average of two months (62/68 mice). Notably, this outcome was independent of any maintenance to the implant – we did not provide solutions to clear the NAPs following implantation nor did we administer any antibiotics. It should also be noted to emphasize the feasibility of this approach that the sole surgeon and *in vivo* experimenter (M.F.R.) was a first-year graduate student upon experimentation who began this project during her rotation phase.

## Discussion

Here we present an indwelling micro-fluidic surgical device which achieves reliable and long-term (up to months) delivery of drugs into the nasal cavity. This represents a tractable solution to a long-standing challenge in pre-clinical drug delivery and discovery – IN delivery. The nasal access port (NAP) we present here allows IN delivery in freely behaving animals as small as mice.

Our findings support multiple uses of the NAP. First, our work indicates that the NAP is suitable for research requiring precise drug delivery to the nasal epithelium, allowing controlled investigation of drug effects, including pharmacokinetics, behavioral responses, and neurochemical/circuitry changes and beyond, relevant to a wide range of research areas, drug discovery, *etc.* The NAP offers an innovative platform for the biomedical community for future testing of the effects of other drugs which are provided IN as even mandated by the FDA in some cases, including candidate analgesics, anti-epileptics, *etc.*, and their neural mechanisms. Second, the NAP proves appropriate for studies needing to examine outcomes of IN substance administration on the body, including bodily organs like the brain, at different scales. By facilitating IN delivery, the NAP bridges the gap between human drug use patterns and preclinical models (which otherwise administer drugs systemically or through jugular catheters – neither of which are used by human drug users). This capability allows for directly translatable research on drug-induced brain changes. Additionally, the small-scale of the NAP affords the ability for simultaneous physiological measures and manipulations. We exemplified that herein by showing the amenability of the NAP for simultaneous cocaine administration during *in vivo* fiber photometry. Additionally, the NAP is indicated when experiments needing to study the outcomes of IN drug/medication on animal behavior. The NAP allows for precise and controlled drug administration in behaving animals, enabling the study of behavioral outcomes of IN drugs. Highly relevant to the study of behavior, the NAP overcomes patency issues that plague traditional jugular catheter use – with the NAP yielding access to the nasal cavity for months without any needed maintenance. Further, animals can freely move and even engage in highly-coordinated instrumental behavioral assays while implanted with the NAP and connected to an infusion system.

There are a few key differences in the application of the NAP for use *in vivo* versus traditional in-dwelling jugular catheters which are the mainstream approach for *in vivo* drug delivery other than manual injections by means of hypodermic needles. First, is that jugular catheters directly access the circulatory system, presenting the risk of infection in that method, and that these catheters must be closed to the environment by a solution which often contains antibiotics, anticoagulants, and glycerol. This maintenance of the jugular catheters, advised by many labs to be performed daily, is not an issue with the NAP since it does not require maintenance. Despite not performing any maintenance on the NAP, in this study we achieved a >91% patency success rate in mice until their pre-determined endpoint. 68 mice remained implanted for 8 weeks, upon which patency was still observed. Also on the topic of patency, while in-dwelling jugular catheter patency is often assayed by monitoring rapid loss of motor tone following injection of propofol or xylazine, both of which are somewhat costly and subject to strict regulations by research institutions and most countries, patency of the NAP can be verified by simple observation of saline or any other innocuous fluid being released from the ipsilateral nostril following a small infusion.

The NAP promises scalability from mice to larger pre-clinical models. We anticipate a comparable approach would achieve reliable and precise drug delivery in rats, rabbits, guinea pigs, pigs, and other pre-clinical models. These adjustments would simply involve 1) providing a greater surface area of the NAP to increase strength of the surgical connection to the nasal bone, 2) increasing the strength of the magnetic attachment configuration or include a latching device to ensure the device stays attached to the nasal bone, and 3) increasing the length of the infusion tube so it can pass through thicker nasal bones. The later adjustment is especially important since where the infusion would occur in the nasal cavity would impact where substances would be distributed – a factor also impacted by intranasal dynamics (*e.g.,* nasal airflow) which may be unique in different animal models^55,56^.

Given the aforementioned advantages stated, of course the NAP requires surgical implantation. This entails aseptic surgical supplies and surgical equipment, making it a potentially costly procedure for a lab not already equipped. The implant also extends beyond the skull-cap, and ultimately for fluid delivery requires a tether – which also may slightly restrict normal head movements which suggests care when interpreting results if applying the NAP for studies on certain behaviors, like orofacial behaviors including facial grooming. While we show proof-of-concept data that the NAP lends itself for use while animals engage in complex instrumental/operant behavioral assays, more work is needed to establish IN self-administration paradigms, including more concretely establishing cocaine-self administration parameters (dose/infusion), and further to study IN self-administration in animals seeking to obtain other psychoactive drugs.

In conclusion, the NAP represents a significant methodology with broad applicability in biomedical and life sciences research, especially in neuroscience, pharmacology, medicinal chemistry, and physiology domains. We anticipate the NAP will be a cornerstone approach for investigating IN drug delivery and its genomic, molecular, cellular, systems, and behavioral consequences – all of which to-date are unexplored. This paradigm promises to allow pre-clinical research and development testing of IN therapies to span beyond *in vitro* models as currently required and closely regulated by both the American Food and Drug Administration (FDA), and European Medicines Agency (EMA)^15^, into whole-animal systems.

## Methods

### Animals

Male and female C57BL/6J mice originating from breeder stock at the Jackson labs (strain #000664; RRID:IMSR_JAX:000664, Bar Harbor, ME), were bred and maintained at the University of Florida. Mice were housed in a temperature-controlled vivarium on a 12:12hr light-dark cycle with *ad libitum* access to food and water. All behavioral testing occurred during the light cycle. Mice were single housed following surgical procedure for chronic implants. Animals were housed in standard shoebox sized plastic cages (Allentown Jaw 75 micro-vent system; Allentown, PA; L: 29.2 cm W: 18.5cm H:12.7cm) in a rack for individually vented cages. Corncob bedding and a Nestlet^TM^ (Ancaes, Bellmore, NY) were in each cage along with manzanita sticks (Bio-Serve) for enrichment. All experimental procedures were conducted within the AALAC animal research program of the University of Florida in accordance with the guidelines from the National Institute of Health and were approved by the University of Florida Institutional Animal Care and Use Committee.

### The nasal access port (NAP)

To achieve microfluidic delivery into the nasal cavity, we developed the nasal access port (NAP). The NAP consists of an infusion port which protrudes 10mm from the Hydex® plastic base which expands 2.8mm (ML) and 9.9mm (AP) to provide sufficient surface area and maximize adhesion to the mouse’s cranium. A 2.54mm magnet is embedded in the base for temporary fastening to a female mating connector for subsequent fluid delivery and/or a dust cap when not in use. Extending 1.3mm beyond the base of the NAP is a 25ga stainless-steel tube (0.5mm OD and 0.029 ID) which is placed inside a small craniotomy over the nasal bone of the animal and permits IN delivery of microliter boluses of fluids with minimal resistance unilaterally in the nasal passage.

### Surgical Procedures

For all surgical procedures, under aseptic conditions, mice were anesthetized with 2-4% Isoflurane in oxygen (IsoFlo®, Patterson Veterinary, Houston, TX) and mounted on a stereotaxic frame while body temperature was maintained on a 38°C heating pad underneath their body. The analgesic Meloxicam (20mg/kg, Patterson Veterinary, Houston, TX) was subcutaneously (s.c.) administered followed by the local anesthetic lidocaine (lidocaine, 3mg/kg, s.c., Patterson Veterinary, Houston, TX) to the scalp before incision and exposure of the skull.

For NAP implantation, the nasal bone was leveled by adjusting the stereotaxic incisor bar. Next, a 1mm diameter hole was drilled through the right nasal bone, 1mm anterior to the frontal/nasal fissure and 0.5mm lateral. These coordinates were selected to position the NAP’s stainless-steel infusion tube immediately upon the vascularized turbinates of the nasal epithelium in mice. Following drilling, the NAP was lowered through the craniotomy, and secured with dental cement (Teets cold cure, Diamond Springs, CA) to the proximal nasal bone and remainder of the skull. After the cement dried, a removable aluminum cap with a magnetic insert was placed over the implant to protect it while not in use.

For *in vivo* monitoring of DA levels, we injected viruses to transduce the GRABDA biosensor. For this, craniotomies were drilled to specifically target the medial shell region of the nucleus accumbens (NAc; +1.25mm AP/0.8mm ML/-4.40mm DV). A pulled micropipette containing 100 nL AAV9.hSyn-GRAB_DA1h (Addgene 113050, titer: 1×10^13^vg/mL)^50^ or 100nL of the inert GRABDA sensor AAV9.hSyn-GRAB_DA-mut (Addgene 140555, titer: 1×10^13^ vg/mL) was slowly lowered into the region of interest and 100nL of viral solution was delivered unilaterally using a Nanoject III (Drummond Scientific, Broomall, PA) at a rate of 2nL/s. Following a 10-minute wait, the pipette was slowly withdrawn from the brain. Subsequently, a 400µm outer diameter, 0.39NA optical fiber terminating in a 2.5mm outer diameter ferrule was lowered to the same coordinates as the injections, and cemented to the skull, for later fiber photometry-based monitoring of DA levels. Animals were allowed to recover for at least 3 weeks prior to any fiber photometry to allow sufficient viral transduction.

Upon completion of surgeries (whether NAP alone or NAP along with methods for monitoring DA levels) the mice were removed from the stereotaxic frame and placed in a clean home cage with on a heating pad to aid recovery. Mice were given Meloxicam (20mg/kg, s.c., Patterson Veterinary, Houston, TX) for a 3-day post-operative period following the surgery.

### Fluid-Delivery Kinetics

We used time-lapse microscopy, using an Olympus epifluorescence microscope (ThorLabs, Newton, NJ), to demonstrate the fast dispensing of small volumes of fluid from the NAP. The access port of the NAP was connected to a female mating connector attached to a line backfilled with 1% fluorescein (Thermo Fisher Scientific), a fluorescent dye, driven by a syringe pump (R-210, Razel Scientific Instruments, Saint Albans, VT) controlled to deliver 1µL aliquots. Time-lapse was acquired using a compact scientific digital camera (CS235CU, ThorLabs, Newton, NJ) and analyzed through the ThorCam software (version 3.7.0.6, ThorLabs, Newton, NJ) across 10 infusion replications. Upon triggering the pump, we were able to validate the fast dispensing of 1µL of the dye with short latency (tens of milliseconds) measured as changes in fluorescence intensity.

### Patency testing

Patency of NAPs was tested a day prior to any behavioral testing occurred. Mice were briefly anesthetized under 2-4% Isoflurane in oxygen and a 1mL syringe attached to a PinPort injector (Instech Laboratories, Plymouth Meeting, PA) was utilized to infuse 0.1mL of saline through the NAP to ensure a small amount of fluid could travel through the NAP and into the nasal cavity as evidenced by a visible drop exiting the ipsilateral nostril. Additionally, to confirm precision and reliability of NAP-mediated IN infusions, a subset of mice were placed in a 15.24 × 15.24 × 15.24 cm (length × width × height) clear acrylic chamber and tethered to a magnetic female connector attached to PU tubing contained within 30.48cm spring connected to a 25ga swivel tethered with the addition of a clean white paper lining the floor of the chamber, and received a 1µL infusion of methylene blue dye. The paper was exchanged for a clean piece and the mice consecutively received 1µL infusions, X4, in a 5s succession. The papers were scanned for visible residues of the dyes. Additionally, a subset of mice underwent NAP-mediated infusions of 1µL, 2µL, or 4µL of methylene blue dye and the nasal cavities were dissected along the midline guided by the frontal incisors for confirmation of ipsilateral infusion site and surface area covered by bolus infusion.

### Cocaine

Cocaine hydrochloride was generously provided by the Drug Discovery Program at the National Institute on Drug Abuse and was dissolved in sterile physiological saline (0.9% NaCl). Concentration was calculated w/v and adjusted to achieve desired concentration per infusion based on body weight of the animal (mg/kg/inf).

### Ultra-performance liquid chromatography-triple quadrupole mass spectrometer (UPLC-MS/MS)

For cocaine and metabolites detection, mice were placed in the clear acrylic chamber and tethered as previously described. In this instance, the line was backfilled with cocaine and attached to a syringe pump triggered by a microcontroller to deliver a single bolus of 8 mg/kg of cocaine (1 µL, X4, every 15s) for tissue collection at 1, 3, 6, and 12 min, post-dose. Following infusion, mice were deeply anesthetized in a chamber filled with isoflurane at T-60 s from the goal time-point. After 45s, lack of response to toe pinch was confirmed and proceeded with tissue harvesting. Blood was collected from cardiac puncture and added to EDTA test tubes containing pre-aliquoted potassium fluoride (Thermo Fisher Scientific) with a final concentration of 2% w/v. Subsequently, blood was centrifuged at 1500 rpm for 10 min at 4°C, the plasma was collected and then stored in -80°C until analysis. Following blood collection, the brain was perfused using 0.5 mL of the same potassium fluoride solution through the heart. The brain was extracted and immediately stored in -80°C for later analysis (**Fig 4**). Plasma and brain samples were stabilized with potassium fluoride to prevent the esterase-mediated metabolism of cocaine.

UPLC-MS/MS was employed to quantify cocaine and its metabolites (benzoylecgonine and ecgonine methyl ester (Millipore Sigma, St. Louis, MO)) in plasma and brain homogenates using a Waters Acquity I-Class system coupled with a Xevo TQ-S Micro triple quadrupole mass spectrometer (Waters, Milford, MA, USA) and running MassLynx (version 4.2, TargetLynx XS; Waters, Milford, MA, USA). Samples were analyzed against freshly prepared calibration and quality control standards in respective matrix following the US-FDA guideline for bioanalytical method validation^57^. These samples were processed via simple protein precipitation method by addition of 80μL of acetonitrile (Thermo Fisher Scientific) containing internal standard (cocaine-*d*3; IS, 2ng/mL, Millipore Sigma St. Louis, MO). Samples were vortex mixed, filtered via a multiscreen Solvinert (Millipore Sigma, St. Louis, MO) 96-well filter plate (0.45μm), centrifuged at 1500 rpm, 4°C for 5 min, and injected onto Waters Acquity UPLC coupled with Xevo® TQ-S micro triple quadruple mass spectrometer. Chromatographic separation was achieved in gradient mode using a Waters Acquity UPLC BEH C18 column (1.7µm, 2.1 x 100mm) with a VanGuard pre-column of a similar chemistry (Waters Co, Milford, MA). The optimized linear gradient program for quantitation of cocaine and its metabolites were as follows: 0-0.8 min, 5% B; 0.8-1.5 min, 5-95% B; 1.5-2.5 min, 95% B; 2.6-3.5 min, 5% B, where mobile phase A was ammonium acetate buffer (2.5 mM; pH: 3.5) and mobile phase B was acetonitrile. The weak needle wash was comprised of water, methanol, and acetonitrile (2:1:1 proportion; 0.1% formic acid). The strong needle wash was comprised of water, acetonitrile, methanol, and isopropyl alcohol (1:1:1:1 proportion; 0.1% formic acid). The column oven and autosampler temperature were set at 50°C and 10°C, respectively. Quantification of cocaine and its metabolites was performed using TargetLynx software. Compound parameters for quantification of analytes using UPLC-MS/MS are represented in **Table 1**.

**Table 1.**
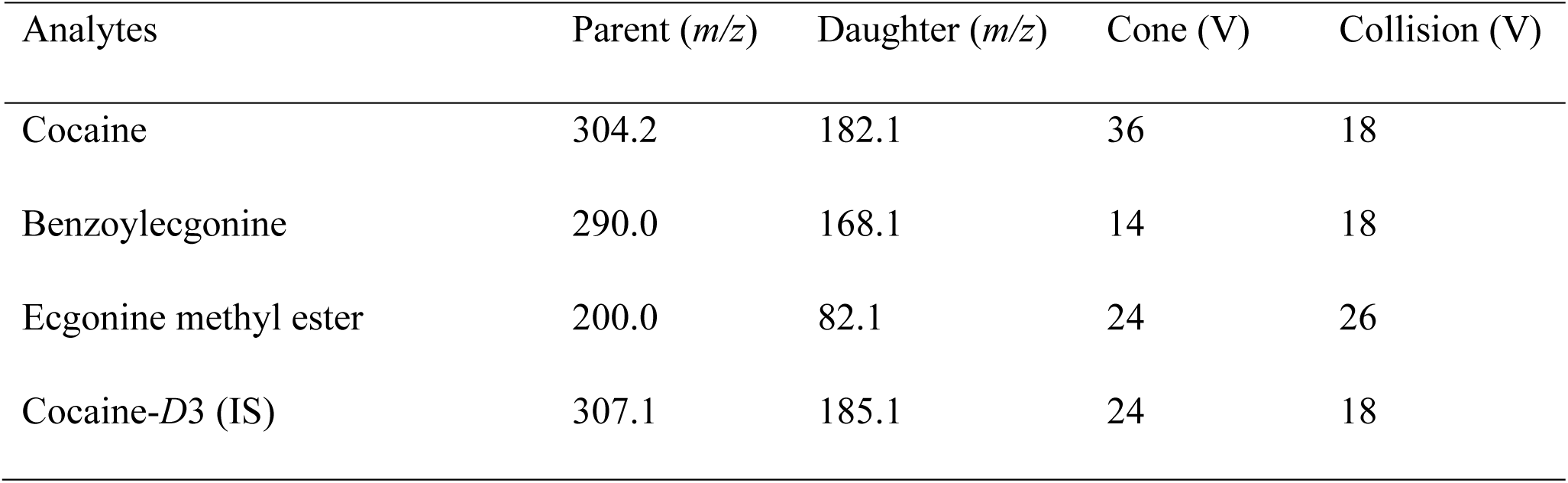
Compound parameters for quantification of analytes using UPLC-MS/MS.

### Behavioral experiments

#### Locomotor activity test for dose response

The locomotor activity test was conducted in a custom-made, open top, acrylonitrile butadiene styrene 30.48 × 30.48 × 30.48cm (length × width × height) dark acrylic chamber. For the dose response, the mice were handled and habituated to the behavior room prior experimentation. On Day 1, the animals were gently scruffed, the protective aluminum cap was removed, and the mice were tethered. The line was empty to allow habituation to the apparatus. On days 2 and 3, baseline locomotion was recorded. The lines were backfilled with sterile saline and the swivel connected to a 1mL syringe in a pump powered and controlled by an Arduino Uno microcontroller running a custom script. The Arduino generated a TTL pulse to trigger a pump to deliver 1µL infusions via the NAP into the nasal epithelium every 15s for a total 4µL bolus. The mice were placed in the chamber for 15 minutes to acclimate each day tethered to the swivel and at the 15 min mark, they received saline infusions and locomotor activity was recorded for 15 minutes post-infusion. At the end of the 15 minutes, mice were gently restrained, the tether was removed, and they were placed back in their home cage. On days 4-15, mice received cocaine hydrochloride in increasing doses daily (0.1mg/kg/inf, 0.3mg/kg/inf, 0.5mg/kg/inf, 1mg/kg/inf, 1.25mg/kg/inf, 1.5mg/kg/inf, 1.75mg/kg/inf, 2mg/kg/inf, 3mg/kg/inf, 4mg/kg/inf, 5mg/kg/inf, 6mg/kg/inf) and followed the same recording schedule previously mentioned. To study the time-course whereby IN cocaine results in hyperlocomotion, a subset of mice used for the locomotor dose-response study above, were delivered eight 1µL infusions (15sec interval) via the NAP into the nasal epithelium of a highly-concentrated cocaine bolus (48mg/kg). All behavioral data were recorded using an infrared video camera (12Hz frame rate) located above each chamber. Locomotion was extracted from the videos off-line in ezTrack^32^.

#### Cocaine Self-Administration

Self-administration studies were conducted in custom-made open-top 60 × 30 × 30 cm (length × width × height) acrylonitrile butadiene styrene (ABS) operant chambers^58^ housed in noise-attenuating cabinets. Each chamber consisted of a guillotine-style door and a wall which contained an extruded panel with 3 nose-poke ports equipped with 880nm infrared photodiodes that generated a TTL pulse when broken to signal port entry. For IN self-administration sessions, a swivel was held by a counterbalanced lever arm anchored to one of the chamber walls. The swivel was connected to a 1mL syringe in a pump and attached to flexible PU tubing contained in a 30.48 cm-long spring which would be attached to the NAP. The operant chambers were powered and driven by an Arduino microcontroller running custom scripts to digitize the occurrence of beam-breaks (nose-pokes, 200Hz sampling rate). Mice underwent 7 days of IN self-administration. Prior to the first session of self-administration, animals were gently scuffed, the protective aluminum cap was removed, and the mice were tethered for 1hr to an empty line to allow habituation to operant chamber. On the next day (session #1), one of the nose-poke ports was randomly assigned as the active port, which would result in cocaine IN infusions, while the other port was deemed as the inactive port and would result in no consequences. This assignment was kept consistent throughout sessions for each mouse. During each 2-hour long session, mice were tethered to the drug line which would deliver 1µL infusion of cocaine (0.3 mg/kg/inf) upon entry of the active nose-poke port. No other cues were present in the chamber, except for the nasal sensation evoked by the delivery of the drug into the nasal epithelium.

#### Fiber photometric recordings of dopamine

Mice were injected into the nucleus accumbens with AAV9.hSyn-GRAB_DA1h^50^ or the inert GRABDA sensor AAV9.hSyn-GRAB_DA-mut and implanted with an optical fiber into the nucleus accumbens shell as described above for later fiber photometry as we have described (see surgical methods)^59^. At least 3 weeks later, 465nm (GRABDA/GRABDA-mut excitation wavelength, driven at 210Hz) and 405nm (UV excitation wavelength, driven at 330Hz; control) light emitting diodes (TDT, RZ10x) were coupled to a fluorescent minicube (Doric lenses, FMC6) using 200µm, 0.22NA FCM optic fiber patch cords. Excitation and emission light were directed through a single 400µm, 0.57NA patch cord connected to the animal by the implanted 2.5mm metal ferrule. Emission light was directed through the mini cube and coupled to femtowatt photoreceivers to monitor GFP and UV signals (TDT, RZ10x). The emission signals were low-pass filtered (6th order), pulse demodulated, and digitized (1kHz) using a Tucker Davis Technologies processor (TDT). All parameters for imaging and acquisition of photometry data were consistent across all groups.

Behavioral sessions for acquiring photometry data lasted approximately 25min. The mice were tethered to the patch cord, the NAP was attached to the infusion line, and then placed in the clear chamber previously mentioned for behavioral monitoring. The animals received saline and cocaine (2mg/kg/inf) intranasally on individual sessions in consecutive days. Each session consisted of a 5 min acclimation period, followed by computer-controlled and user-initiated infusions via a pump. Each bolus cycle consisted of 1µL aliquots delivered every 15s, X4 for a total volume of 4µL bolus. The session consisted of 3 bolus repetitions every 5 minutes to allow monitoring of DA transients.

### Histology

At the end of the experiment, mice were overdosed with sodium pentobarbital (Fatal-Plus, Patterson) and transcardially perfused with cold 0.9% saline followed by 10% phosphate-buffered formalin (SF100-4, Thermo Fisher Scientific). Brains were removed and stored at 4°C in 30% sucrose formalin for cryoprotection and postfixation for at least 24 h before they were coronally sectioned at 40 µm on a freezing microtome (Leica). Sections were slide mounted and coverslipped with a 4′,6-diamidino-2-phenylindole (DAPI)-counterstained mounting medium (DAPI-fluoromount-G, SouthernBiotech, Birmingham, AL). Location of the nucleus accumbens, the target region for fiber photometry, was identified utilizing a mouse brain atlas while referring the DAPI label^60^. Viral injections and optical fiber placements were verified on a Nikon Ti2e inverted fluorescent microscope and only the mice with on target injections/fiber implants contributing to the data set.

### Statistical Analysis

All data were processed using semi-automated steps whenever possible, including processing data from mixed treatment groups at the same time. Statistical comparisons were analyzed in GraphPad Prism which included repeated measures (rm) and/or two-way ANOVAs followed by Tukey’s multiple comparison corrections when needed. All data are reported as mean±SEM unless noted otherwise.

## Data availability

The authors will provide original full data upon reasonable requests.

## Code availability

No custom code was used in this study.

## Acknowledgements

We thank Ellyse Thomas for expert animal care assistance and Intech laboratories for working with the authors (M.F.R and D.W.W.) to manufacturer the NAP. This work was supported by R01DA049449 and funds from the University of Florida to D.W.W.. M.F.R is supported by T3DC2015994.

## Author Contributions

Conceptualization: MFR & DWW Methodology: MFR, AG, AS, SES, DWW Investigation: MFR, EW, SES, AG, DWW Visualization: MFR & DWW

Funding acquisition: MFR & DWW Supervision: DWW

Writing – original draft: MFR & DWW

Writing – review & editing: MFR, EW, AG, AS, SES, DWW

## References

1. Bousquet, J. et al. Allergic rhinitis. Nat. Rev. Dis. Prim. 6, 95 (2020).

2. Serra, S. et al. Intranasal Fentanyl for Acute Pain Management in Children, Adults and Elderly Patients in the Prehospital Emergency Service and in the Emergency Department: A Systematic Review. J. Clin. Med. 12, (2023).

3. Smith-Apeldoorn, S. Y., Veraart, J. K., Spijker, J., Kamphuis, J. & Schoevers, R. A. Maintenance ketamine treatment for depression: a systematic review of efficacy, safety, and tolerability. The lancet. Psychiatry 9, 907–921 (2022).

4. Kehagia, E., Papakyriakopoulou, P. & Valsami, G. Advances in intranasal vaccine delivery: A promising non-invasive route of immunization. Vaccine 41, 3589–3603 (2023).

5. Lipton, R. B. et al. Safety, tolerability, and efficacy of zavegepant 10 mg nasal spray for the acute treatment of migraine in the USA: a phase 3, double-blind, randomised, placebo-controlled multicentre trial. Lancet. Neurol. 22, 209–217 (2023).

6. Sharma, S. & Detyniecki, K. Rescue therapies in epilepsy. Curr. Opin. Neurol. 35, 155– 160 (2022).

7. 7. NIDA. NIDA Commonly Used Drugs Charts. https://nida.nih.gov/research-topics/commonly-used-drugs-charts.

8. Kiluk, B. D., Babuscio, T. A., Nich, C. & Carroll, K. M. Smokers versus snorters: do treatment outcomes differ according to route of cocaine administration? Exp. Clin. Psychopharmacol. 21, 490–498 (2013).

9. Molteni, M., Saibene, A. M., Luciano, K. & Maccari, A. Snorting the clivus away: an extreme case of cocaine-induced midline destructive lesion. BMJ Case Rep. 2016, (2016).

10. Messinger, E. Narcotic septal perforations due to drug addiction. JAMA 179, 964–965 (1962).

11. Goodger, N. M., Wang, J. & Pogrel, M. A. Palatal and nasal necrosis resulting from cocaine misuse. Br. Dent. J. 198, 333–334 (2005).

12. Peyrière, H. et al. Necrosis of the intranasal structures and soft palate as a result of heroin snorting: a case series. Subst. Abus. 34, 409–414 (2013).

13. Costantino, H. R., Illum, L., Brandt, G., Johnson, P. H. & Quay, S. C. Intranasal delivery: physicochemical and therapeutic aspects. Int. J. Pharm. 337, 1–24 (2007).

14. Chung, S. et al. The nose has it: Opportunities and challenges for intranasal drug administration for neurologic conditions including seizure clusters. Epilepsy Behav. Reports 21, 100581 (2023).

15. Trows, S., Wuchner, K., Spycher, R. & Steckel, H. Analytical Challenges and Regulatory Requirements for Nasal Drug Products in Europe and the U.S. Pharmaceutics vol. 6 195– 219 at 10.3390/pharmaceutics6020195 (2014).

16. Huang, H. et al. Chronic and acute intranasal oxytocin produce divergent social effects in mice. Neuropsychopharmacology 39, 1102–1114 (2014).

17. Hanson, L. R., Fine, J. M., Svitak, A. L. & Faltesek, K. A. Intranasal administration of CNS therapeutics to awake mice. J. Vis. Exp. (2013) doi:10.3791/4440.

18. Gothwal, A., Lamptey, R. N. L. & Singh, J. Multifunctionalized Cationic Chitosan Polymeric Micelles Polyplexed with pVGF for Noninvasive Delivery to the Mouse Brain through the Intranasal Route for Developing Therapeutics for Alzheimer’s Disease. Mol. Pharm. 20, 3009–3019 (2023).

19. Rojanaratha, T. et al. Preparation, physicochemical characterization, ex vivo, and in vivo evaluations of asiatic acid-loaded solid lipid nanoparticles formulated with natural waxes for nose-to-brain delivery. Eur. J. Pharm. Sci. 203, 106935 (2024).

20. Wesson, D. W., Donahou, T. N., Johnson, M. O. & Wachowiak, M. Sniffing behavior of mice during performance in odor-guided tasks. Chem Senses 33, 581–596 (2008).

21. Verhagen, J. V, Wesson, D. W., Netoff, T. I., White, J. A. & Wachowiak, M. Sniffing controls an adaptive filter of sensory input to the olfactory bulb. Nat Neurosci 10, 631– 639 (2007).

22. 22. Gulner, B. R., Navabi, Z. S. & Kodandaramaiah, S. B. 3D morphometric analysis of mouse skulls using microcomputed tomography and computer vision. bioRxiv 2022.10.26.513830 (2022) doi:10.1101/2022.10.26.513830.

23. Gunaydin, L. A. et al. Natural Neural Projection Dynamics Underlying Social Behavior. Cell 157, 1535–1551 (2014).

24. Benuck, M., Lajtha, A. & Reith, M. E. Pharmacokinetics of systemically administered cocaine and locomotor stimulation in mice. J. Pharmacol. Exp. Ther. 243, 144–9 (1987).

25. Shang, L. et al. Catalytic activities of a highly efficient cocaine hydrolase for hydrolysis of biologically active cocaine metabolites norcocaine and benzoylecgonine. Sci. Rep. 13, 640 (2023).

26. Stewart, D. J., Inaba, T., Lucassen, M. & Kalow, W. Cocaine metabolism: cocaine and norcocaine hydrolysis by liver and serum esterases. Clin. Pharmacol. Ther. 25, 464–8 (1979).

27. Spiehler, V. R. & Reed, D. Brain concentrations of cocaine and benzoylecgonine in fatal cases. J. Forensic Sci. 30, 1003–11 (1985).

28. Volkow, N. D., Wise, R. A. & Baler, R. The dopamine motive system: implications for drug and food addiction. Nat. Rev. Neurosci. 18, 741–752 (2017).

29. Carelli, R. M. Nucleus accumbens cell firing and rapid dopamine signaling during goal-directed behaviors in rats. Neuropharmacology 47, **Supple**, 180–189 (2004).

30. Cornish, J. L. & Kalivas, P. W. Repeated cocaine administration into the rat ventral tegmental area produces behavioral sensitization to a systemic cocaine challenge. Behav. Brain Res. 126, 205–209 (2001).

31. Sun, F. et al. A Genetically Encoded Fluorescent Sensor Enables Rapid and Specific Detection of Dopamine in Flies, Fish, and Mice. Cell 174, 481–496.e19 (2018).

32. Pennington, Z. T. et al. ezTrack: An open-source video analysis pipeline for the investigation of animal behavior. Sci. Rep. 9, 19979 (2019).

33. Carelli, R. M., King, V. C., Hampson, R. E. & Deadwyler, S. A. Firing patterns of nucleus accumbens neurons during cocaine self-administration in rats. Brain Res. 626, 14–22 (1993).

34. Stuber, G. D., Roitman, M. F., Phillips, P. E. M., Carelli, R. M. & Wightman, R. M. Rapid Dopamine Signaling in the Nucleus Accumbens during Contingent and Noncontingent Cocaine Administration. Neuropsychopharmacology 30, 853–863 (2005).

35. Mu, P. et al. Exposure to Cocaine Dynamically Regulates the Intrinsic Membrane Excitability of Nucleus Accumbens Neurons. J. Neurosci. 30, 3689–3699 (2010).

36. Suska, A., Lee, B. R., Huang, Y. H., Dong, Y. & Schluter, O. M. Selective presynaptic enhancement of the prefrontal cortex to nucleus accumbens pathway by cocaine. Proc. Natl. Acad. Sci. 110, 713–718 (2013).

37. Piazza, P. V, Deroche-Gamonent, V., Rouge-Pont, F. & Le Moal, M. Vertical shifts in self-administration dose-response functions predict a drug-vulnerable phenotype predisposed to addiction. J. Neurosci. 20, 4226–32 (2000).

38. Piazza, P. V, Deminière, J. M., Le Moal, M. & Simon, H. Factors that predict individual vulnerability to amphetamine self-administration. Science 245, 1511–3 (1989).

39. Koob, G. F. & Moal, M. Le. Drug Abuse: Hedonic Homeostatic Dysregulation. Science (80-.). 278, 52–58 (1997).

40. Kreek, M. J. & Koob, G. F. Drug dependence: stress and dysregulation of brain reward pathways. Drug Alcohol Depend. 51, 23–47 (1998).

41. Deminiere, J. M., Piazza, P. V., Le Moal, M. & Simon, H. Experimental approach to individual vulnerability to psychostimulant addiction. Neurosci. Biobehav. Rev. 13, 141– 147 (1989).

42. Stretch, R., Gerber, G. J. & Wood, S. M. Factors Affecting Behavior Maintained by Response-Contingent Intravenous Infusions of Amphetamine in Squirrel Monkeys. Can. J. Physiol. Pharmacol. 49, 581–589 (1971).

43. de Wit, H. & Stewart, J. Reinstatement of cocaine-reinforced responding in the rat. Psychopharmacology (Berl). 75, 134–43 (1981).

44. Shaham, Y., Shalev, U., Lu, L., de Wit, H. & Stewart, J. The reinstatement model of drug relapse: history, methodology and major findings. Psychopharmacology (Berl). 168, 3–20 (2003).

45. Katz, J. L. & Higgins, S. T. The validity of the reinstatement model of craving and relapse to drug use. Psychopharmacology (Berl). 168, 21–30 (2003).

46. Kasanetz, F. et al. Transition to addiction is associated with a persistent impairment in synaptic plasticity. Science 328, 1709–12 (2010).

47. Deroche-Gamonet, V., Belin, D. & Piazza, P. V. Evidence for Addiction-like Behavior in the Rat. Science (80-. ). 305, 1014–1017 (2004).

48. Grahame, N. J., Phillips, T. J., Burkhart-Kasch, S. & Cunningham, C. L. Intravenous cocaine self-administration in the C57BL/6J mouse. Pharmacol. Biochem. Behav. 51, 827–834 (1995).

49. Kuzmin, A. Reinforcing and Neurochemical Effects of Cocaine Differences Among C57, DBA, and 129 Mice. Pharmacol. Biochem. Behav. 65, 399–406 (2000).

50. Griffin, W. C. & Middaugh, L. D. Acquisition of lever pressing for cocaine in C57BL/6J mice: effects of prior Pavlovian conditioning. Pharmacol. Biochem. Behav. 76, 543–549 (2003).

51. Slosky, L. M. et al. Establishment of multi-stage intravenous self-administration paradigms in mice. Sci. Rep. 12, 21422 (2022).

52. Thomsen, M. & Caine, S. B. Intravenous Drug Self-administration in Mice: Practical Considerations. Behav. Genet. 37, 101–118 (2007).

53. Li, Y. et al. The optimized jugular vein catheterization reinforced cocaine self-administration addictive model for adult male Sprague–Dawley rats. Sci. Rep. 12, 11711 (2022).

54. Thomsen, M. & Caine, S. B. Chronic intravenous drug self-administration in rats and mice. Curr. Protoc. Neurosci. Chapter 9, Unit 9.20 (2005).

55. Morgan, K. T. & Monticello, T. M. Airflow, gas deposition, and lesion distribution in the nasal passages. Environ. Health Perspect. 85, 209–218 (1990).

56. Wu, Z., Jiang, J., Lischka, F. W. & Zhao, K. Is the mouse nose a miniature version of a rat nose? A computational comparative study. Comput. Methods Programs Biomed. 254, 108282 (2024).

57. 57. Bioanalytical Method Validation and Study Sample Analysis. Center for Drug Evaluation and Research Center for Biologics Evaluation and Research https://www.fda.gov/regulatory-information/search-fda-guidance-documents/m10-bioanalytical-method-validation-and-study-sample-analysis (2022).

58. Wright, K. N. & Wesson, D. W. The tubular striatum and nucleus accumbens distinctly represent reward-taking and seeking. J. Neurophysiol. 125, 166–183 (2021).

59. Johnson, N. L. et al. Dopaminergic signaling to ventral striatum neurons initiates sniffing behavior. Nat. Commun. 16, (2025).

60. Paxinos, G. & Franklin, K. The Mouse Brain in Stereotaxic Coordinates. (Academic Press, San Diego, 2000).

